# Function expansion of antitumor transcriptional activator NFE2L1 by the original discovery of its non-transcription factor activity

**DOI:** 10.1101/2020.10.08.330597

**Authors:** Qiu Lu, Yang Qiufang, Li Peng, Zhou Xiaowen, Yang Yonghui, Zhou Xiuman, Gou Shanshan, Zhai Wenjie, Li Guodong, Ren Yonggang, Zhao Wenshan, Wu Yahong, Qi Yuanming, Gao Yanfeng

## Abstract

Antitumor transcription activator NFE2L1, with the functions to regulate redox homeostasis, protein turnover, and material metabolism, plays an important role in embryonic development and specialization of tissue and organ functions. Deficiency of *NFE2L1* gene in different regions yields distinct phenotypes, suggesting that NFE2L1 may have a transcription factor-independent function. Here we originally discovered the non-transcription factor activity of NFE2L1 by constructing a truncated protein-NFE2L1^ΔC^ without 152 aa at the C-terminus which lost the transcription factor activity. The regulation of NFE2L1 on redox homeostasis, proteasome function, and immune response mainly depends on its transcription activator function in nucleus, while the regulation on metabolism, ribosome function, and canceration is germanely to its non-transcription factor activity in cytoplasm. Surprisingly, the results indicated the tumor suppressive effect of NFE2L1 by repression of Wnt/β-catenin signaling in a non-transcription factor manner, indicating the potential value of NFE2L1 as a therapeutic target in clinical cancer treatment independent of its transcription factor activity. Our observations reveal the non-transcription factor activity of NFE2L1 for the first time, and lay foundation for the basic and applied research of NFE2L1.

## INTRODUCTION

Redox homeostasis which composed of oxidative stress and antioxidant systems is a necessary prerequisite for maintaining normal physiological homeostasis. The long-term oxidative damage caused by the imbalance of redox homeostasis is the main reason for various chronic systemic diseases, such as chronic inflammation, diabetes, obesity, neurodegenerative diseases, aging, tumors, etc[1–3]. The cap’n’collar basic-leucine zipper (CNC-bZIP) transcription factor family is the most powerful transcription factor group in eukaryotes involved in regulating intracellular redox homeostasis. Among the 6 members of the mammalian CNC-bZIP transcription factor (Fig. 1A), the nuclear factor erythroid 2 like 1 (NFE2L1/Nrf1) contains the most protein subtypes with complicated regulation functions. Also, NFE2L1/Nrf1 is the only member in this family whose knock-out leads to embryonic lethality[2].

**Fig. 1.**
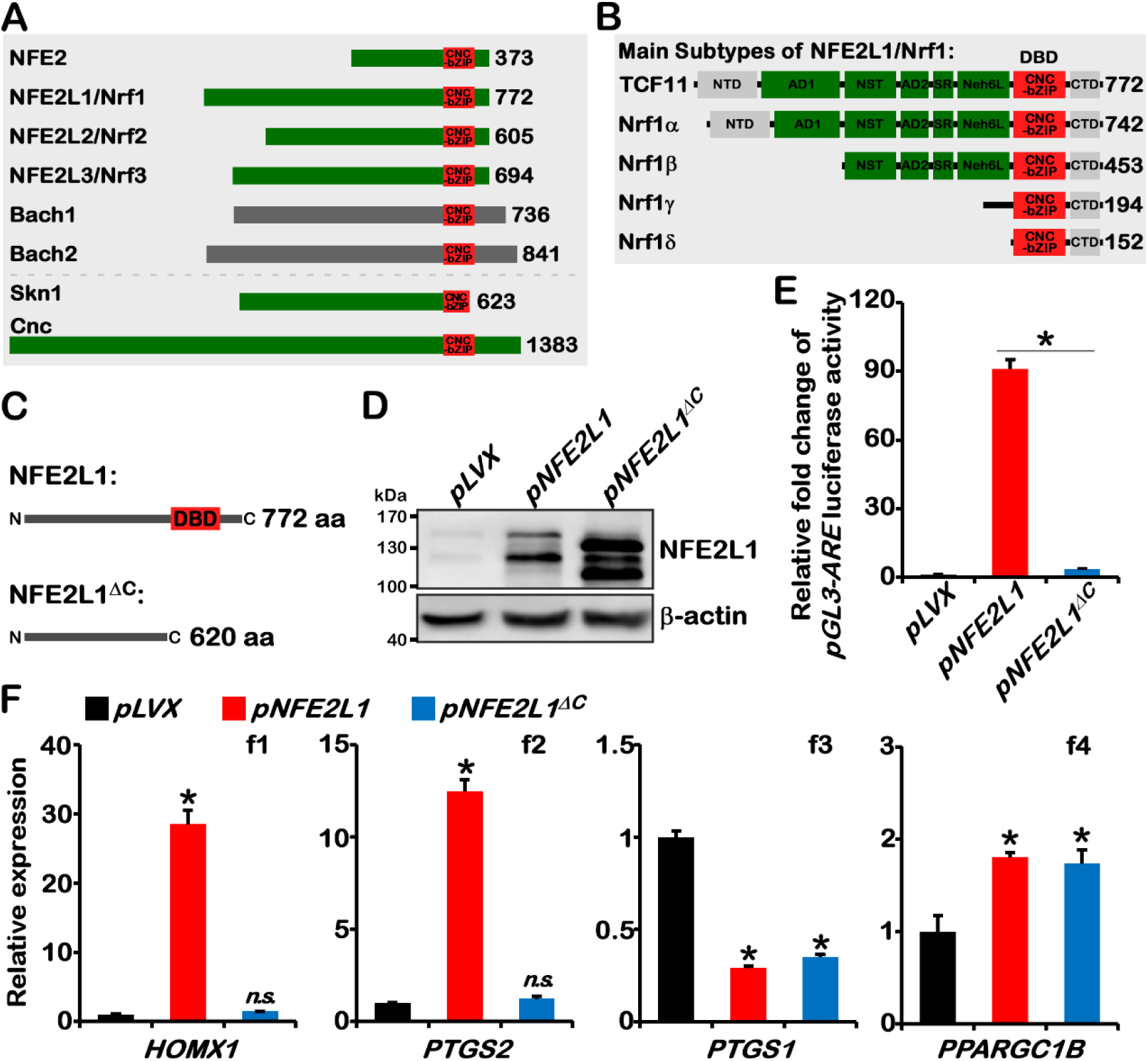
NFE2L1’s regulation of downstream genes is not entirely dependent on its transcription factor function. (A) Members of the CNC-bZIP transcription factor family: NFE2, NFE2L1, NFE2L2, NFE2L3, BACH1, BACH2; Skn1 is the homologous protein in *Caenorhabditis elegans*; Cnc is the homologous protein in *Drosophila melanogaster*. (B) There are five main subtypes (TCF11 (772 aa), Nrf1α (742 aa), Nrf1β (453 aa), Nrf1γ (194 aa), Nrf1δ (152 aa)) and eight main functional domains (NTD, AD1, NST, AD2, SR, Neh6L, DBD, CTD) of NFE2L1. (C) Diagram of NFE2L1^ΔC^: 152 amino acid residues are deleted at the C-terminus, and the DBD structure is lost. (D) The *pLVX*, *pNFE2L1*, *pNFE2L1*^*ΔC*^ plasmids were transfected into HepG2 cells respectively (concentration: 2 μg/mL), and the total protein was collected after 24 hours, then the expression of NFE2L1 and NFE2L1^ΔC^ were detected by WB. (E) The *pGL3-ARE* and the internal reference plasmid *pRL-TK* were transfected into HepG2 cells with *pLVX*, *pNFE2L1*, and *pNFE2L1*^*ΔC*^ respectively, and the dual luciferase activity was detected after 24 hours. n ≥ 3, ‘**\***’ means *p* < 0.05. (F) The *pLVX*, *pNFE2L1*, *pNFE2L1*^*ΔC*^ plasmids were transfected into HepG2 cells respectively, the total RNA was collected and reverse transcribed after 24 h, and the expression of *HOMX1*, *PTGS2*, *PTGS1*, *PPARGC1B* were detected by qPCR, with the *β-actin* as the internal reference gene. n ≥ 3, ‘**\***’ means compared with *pLVX* group and *p* < 0.05, ‘*n.s.*’ means ‘No Significance’ compared with *pLVX* group.

Homo sapiens *NFE2L1* gene which contains 7 exons and 23 transcripts was cloned in 1993[4]. The five main subtypes of NFE2L1 protein are TCF11 (~120-140kDa)[5], Nrf1α (~120-140kDa)[4], Nrf1β (~65kDa)[6], Nrf1γ (~36kDa), and Nrf1δ (~25kDa)[7] (Fig. 1B). According to the structure and functional characteristics [2], NFE2L1 protein has been divided into eight main functional structural regions: N-terminal domain (NTD, aa 1-124), acidic domain 1 (AD1, aa 125-328), asparagine/serine/threonine region (NST, aa 329-433), acidic domain 2 (AD2, aa 434-482), serine-repeat region (SR, aa 483-518), NFE2L2-ECH homology 6 like region (Neh6L, aa 519-610), DNA binding domain (DBD, aa 611-716), and C-terminal domain (CTD, aa 717-772). The NFE2L1 protein in the cytoplasm binds to the endoplasmic reticulum (ER) membrane through the NTD, Neh6L, and CTD domains, and places the AD1, NST, AD2, and SR regions between NTD and Neh6L in the ER lumen; while the DBD domain between Neh6L and CTD is located in the cytosol[2].

*NFE2L1*, which widely expresses in human tissues, is sensitive to environmental changes. Oxidative stimulation triggered by drugs, heavy metals, radiation, nutrition, metabolites, and even changes in ambient temperature results in differential expression of the *NFE2L1* gene[2,8]. The molecular function of NFE2L1 involves the regulation of intracellular redox homeostasis, carbohydrate and lipid metabolism, protein turnover, while the physiological function of NFE2L1 contains antioxidant detoxification, embryonic development, organ function, canceration, inflammation, and immune responses[2,8]. In particular, *NFE2L1* displays a strong inhibitory effect on human-derived tumor cell lines, and the specific knockout of *NFE2L1* in mouse liver triggers the occurrence of hepatocellular carcinoma (HCC) without exogenous stimulation[9–11]. Recent studies reveal that NFE2L1, a crucial transcription factor for neonatal heart regeneration, can promote cardiomyocyte survival[12]. The role of *NFE2L1* in chronic inflammation, cancer, diabetes, obesity, neurodegenerative diseases and other chronic systemic diseases suggests that *NFE2L1* has potentials in clinical application.

Our previous studies revealed the function of various subtypes and domains and the potential anti-tumor effect of NFE2L1. In addition, we found that the “NFE2L1 paradox”, which overexpressing or knocking out *NFE2L1* causes the same trend of changes in downstream genes, was caused by the inhibition of NFE2L2 by NFE2L1[7,10,11,13–15]. However, there are still many unsolved problems in the study of NFE2L1. For example, mice with different regions of the *NFE2L1* gene knocking out possess different phenotypes[16–18]. NFE2L1 and NFE2L2 share the same antioxidant/electrophile-response element (ARE/EpRE) but exhibit different functions [19–22]; Some downstream genes are repressed by NFE2L1[23], and part of downstream genes have no ARE element in their promoter region[10]. These questions imply that the function of NFE2L1 may not only depend on its transcription activator activity. However, there are no reports of the possible non-transcription factor functions of NFE2L1, and the existing modification-cleavage theory and multiple protein subtype models are all based on the transcription factor function of NFE2L1. In this study, we originally explored the non-transcription factor function of NFE2L1.

## RESULTS

### NFE2L1^ΔC^ which lacks the DBD structure and has no transcription factor activity can regulate partial of downstream genes

The DBD domain, containing CNC and bZIP, is the structural basis for NFE2L1 to form the heterodimer, bind to DNA, and enter into the nucleus[24,25]. In order to verify whether NFE2L1 has functions other than transcription activators, an expression plasmid was constructed to express the truncated protein-NFE2L1^ΔC^ which lacking the DBD and CTD domains (Fig. 1C). The results showed that the NFE2L1^ΔC^ with the C-terminal 152 aa deleted in the SDS-PAGE gel was about 10 kDa smaller than the full-length NFE2L1 by Western blot (WB) (Fig. 1D). Moreover, prolifically NFE2L1^ΔC^ were detected, with the same amount of plasmids transfected, indicating that the NFE2L1 protein was more stable after the C-terminal 152 aa was deleted. And these results were consistent with previous results reported by Zhang's finding (other labs) that CTD deletion activated the NFE2L1’s transcription factor function[7].

After the NFE2L1^ΔC^ protein was successfully expressed, transcription factor activity of the NFE2L1^ΔC^ protein was tested by the dual luciferase activity report experiment and the results indicated that its transcriptional activation activity on ARE was significantly reduced despite the high protein abundance of NFE2L1^ΔC^ (Fig. 1E). Subsequently, the response of some classic downstream genes to NFE2L1 and NFE2L1^ΔC^ was tested, and *HOMX1*[26], and *PTGS2*[10,27] were not transcribed by NFE2L1^ΔC^ (Fig. 1F), while *PTGS1*[10], and *PPARGC1B*[28] could still respond to NFE2L1^ΔC^ without transcription factor activity (Fig. 1F). These results strongly proved that NFE2L1 had other functions independent of its transcription factor activity.

### Transcriptome sequencing highlighted the non-transcription factor activity of NFE2L1

To further verify and clarify the non-transcription factor function of NFE2L1, transcriptome sequencing of NFE2L1 and NFE2L1^ΔC^ stable-overexpressing cell lines were constructed using the lentiviral system. The principal component analysis (PCA) results demonstrated that nine sequenced samples were clearly divided into three categories and consistent with their respective groupings (Fig. 2A). The heatmap of distance and the cluster tree both indicated that overexpression of NFE2L1 and NFE2L1^ΔC^ caused changes in the transcriptome (Fig. 2B, 2C).

**Fig. 2.**
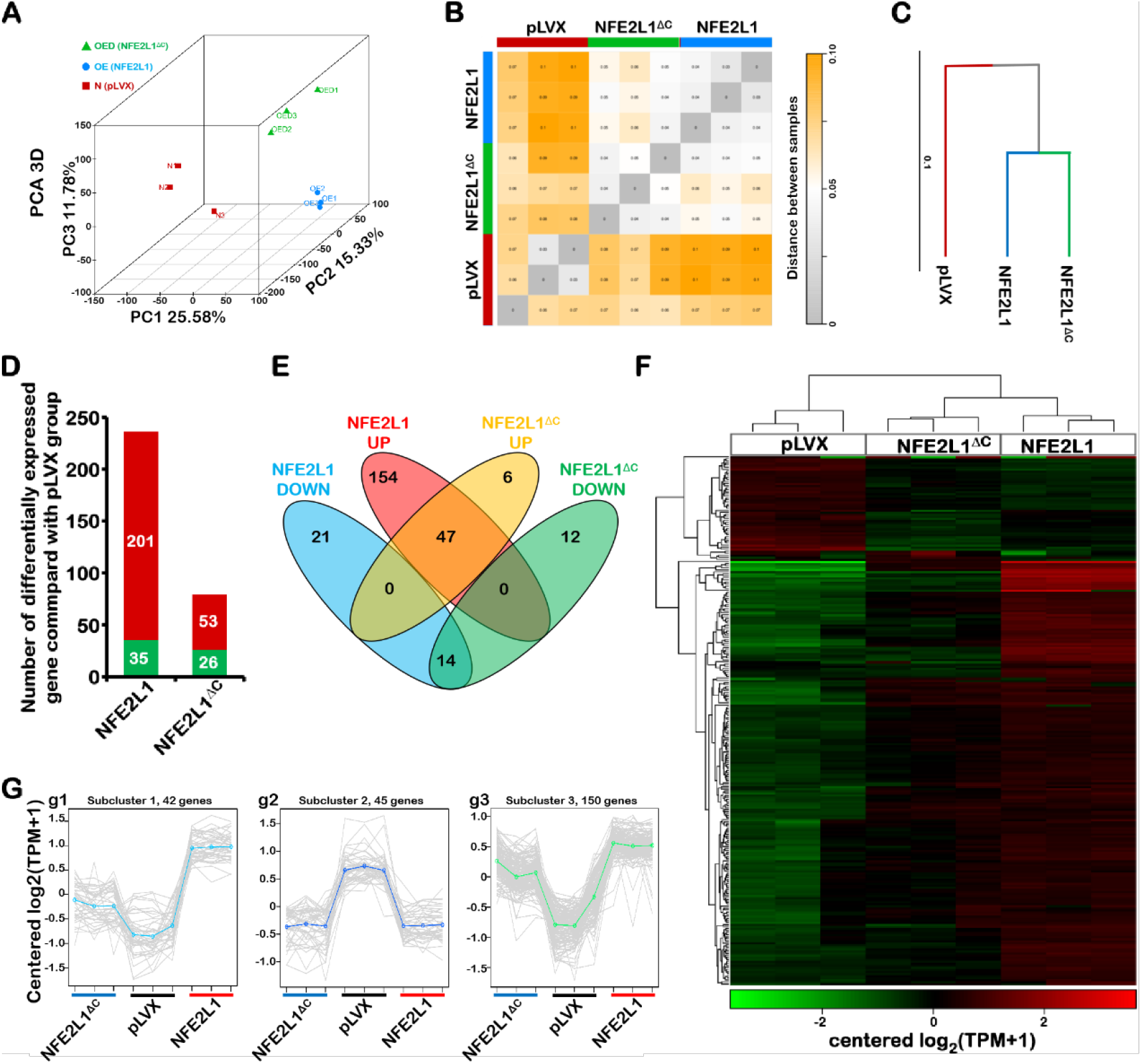
NFE2L1 and NFE2L1^ΔC^ have different regulation patterns on downstream genes. (A) PCA result of component 1, component 2 and component 3 for 9 samples. The closer the distance, the higher the similarity. (B) Heatmap of distance values across samples. The smaller the value (the grayer the color block), the higher the similarity between samples; the larger the value (the yellower the color block), the farther the distance. (C) Cluster tree shows the distance between three groups of samples. The length of the branches represents the distance between the samples, the more similar the samples the closer they are. (D) The number of DEGs in the NFE2L1 and NFE2L1^ΔC^ groups (compared to the pLVX group, respectively), the ‘red band’ indicates increased expression, and the ‘green band’ indicates decreased expression. (E) Venn diagrams display the association of DEGs in NFE2L1 and NFE2L1^ΔC^ groups. The heatmap (F) and gene expression patterns (G) shows the clustering of DEGs in pLVX, NFE2L1, NFEL1^ΔC^ groups.

Then, the conditions for the Transcripts Per Million (TPM) value and the expression fold change of the gene (TPM > 5, |Fold Change| > 2, *p*-Value < 0.01) was set to screen out the differentially expressed genes (DEGs) that caused by stable overexpression of NFE2L1 or NFE2L1^ΔC^. According to the change in the number of DEGs, it could be found that the number of up-regulated genes reduced from 201 to 53 (73.6% decrease) and the number of down-regulated genes reduced from 35 to 26 (25.7% decrease) with NFE2L1 lost the function of transcription factors (Fig. 2D). Venn diagram analysis of DEGs demonstrated that 88.7% (47/53) of the upregulated DEGs in the NFE2L1^ΔC^ group were consistent with the DEGs of the NFE2L1 group and the ratio of the down-regulated DEGs was 53.8% (14/26) (Fig. 2E).

From the heatmap clustering and expression pattern classification, it had been shown that DEGs are mainly divided into three categories. The first category: the genes could be activated by NFE2L1 instead of NFE2L1^ΔC^; the second category: the genes were suppressed by NFE2L1 and NFE2L1^ΔC^; the third category: genes could be activated both by NFE2L1 and NFE2L1^ΔC^ (Fig. 2F, 2G). Based on the analysis of DEGs, it could be found that NFE2L1’s transcriptional inhibition of downstream genes are more relevant to its non-transcription factor function.

### NFE2L1^ΔC^ cannot affect the activity of proteasome

Ubiquitination-proteasome system (UPS), which degrades ubiquitination-modified proteins and participates in many important physiological and biochemical processes such as growth and differentiation of cells, replication and repair of DNA, metabolism of substances and energy, and immune responses, is the main protein degradation system in cells [29]. The transcriptional regulation of the proteasome subunit gene (*PSMs*) is one of the most classical transcription factor function of NFE2L1[30,31]. Although transcriptome sequencing results showed that NFE2L1 possessed activity independent of its transcription factor function, but the NFE2L1^ΔC^, theoretically lost the transcription factor activity, should have no effects on the proteasome. Next, this inference was verified in the pLVX, NFE2L1 and NFE2L1^ΔC^ cell lines. The ubiquitination level of the total protein was decreased by the overexpression of NFE2L1, which meant that the function of proteasome was enhanced, while this phenomenon had not been observed after NFE2L1^ΔC^ overexpression (Fig. 3A). The results of heat map analysis (Fig. 3B) and total expression (Fig. 3C) of *PSMs* in the three cell lines showed that *PSMs* responded to NFE2L1 instead of NFE2L1^ΔC^, which was consistent with the changes in total protein ubiquitination. The expression of *PSMs* was significantly up-regulated due to the overexpression of NFE2L1 and the fold change was mostly less than twice (Fig. 3D). The results were in line with our earlier findings that NFE2L1 moderately regulated the transcription of *PSMs* without exogenous stimulation[15].

**Fig. 3.**
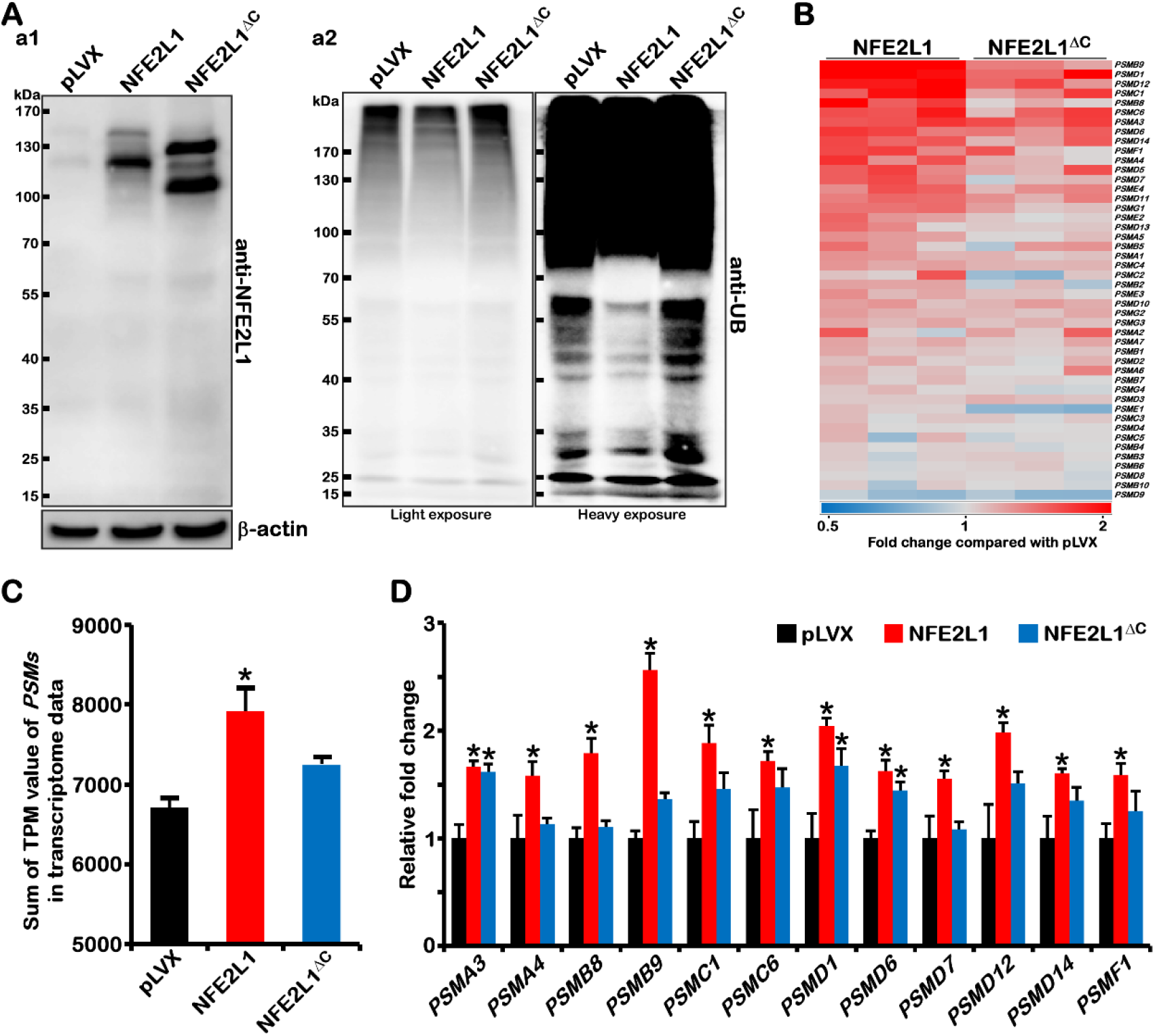
The effect of NFE2L1 and NFE2L1^ΔC^ on proteasome function. (A) Extract the total protein of pLVX, NFE2L1, NFEL1^ΔC^ cell lines, the expression of NFE2L1 protein (a1) and the ubiquitination of total protein (a2: light exposure results on the left, heavy exposure results on the right) were detected by WB. (B) The heat map shows the expression of *PSMs* after overexpression of NFE2L1 or NFEL1^ΔC^ (red indicates an increase, blue indicates a decrease, and the data comes from transcriptome sequencing). (C) The sum of the TPM values of *PSMs* in pLVX, NFE2L1, NFEL1^ΔC^ cell lines. (D) The histogram shows the relative fold change of some *PSMs* with significant differences after overexpression of NFE2L1or NFEL1^ΔC^ protein. n = 3, ‘**\***’ means compared with pLVX group and *p* < 0.05.

### Distinguishing the function of NFE2L1’s transcription activator function and non-transcription factor activity

To further analyze the functional difference between NFE2L1 and NFE2L1^ΔC^, the screening conditions (TPM>0.1, |Fold Change|>1.5, *p*-Value<0.01) were expanded to obtain more DEGs (Fig. S1A). 137 downstream genes (123 up-regulated and 14 down-regulated) which were regulated by NFE2L1’s transcription activator function, not affected by NFE2L1^ΔC^ (Fig. S1B, Table S1). And 262 DEGs (131 up-regulated and 131 down-regulated) that related to NFE2L1’s non-transcription factor activity been acquired, which comes from the intersection of NFE2L1/pLVX group with NFE2L1^ΔC^/pLVX group and have no significant change in NFE2L1/ NFE2L1^ΔC^ group (Fig. S1C, Table S2).

Then, functional cluster analysis on these DEGs were performed to distinguish the transcription factor function and non-transcription factor function of NFE2L1.There were twenty KEGG pathways with *p*-value < 0.01 obtained from the enrichment of DEGs from the NFE2L1/pLVX group. Among them, ten pathways have been involved in infection, inflammation and immunity (Measles, Herpes simplex infection, Influenza A, Epstein-Barr virus infection, Legionellosis, Viral carcinogenesis, TNF signaling pathway, Hepatitis C, Hepatitis B, HTLV-I infection). Eight pathways participated in development, cancer, and canceration (p53 signaling pathway, PI3K-Akt signaling pathway, Pathways in cancer, Wnt signaling pathway, Viral carcinogenesis, TNF signaling pathway, Breast cancer, Hippo signaling pathway). The other two pathways have been implicated in metabolism (Phenylalanine metabolism, Regulation of lipolysis in adipocyte), and the other two are involved in protein turnover (Proteasome, Ribosome biogenesis in eukaryotes) (Figs. 4A and S1D).

**Fig. 4.**
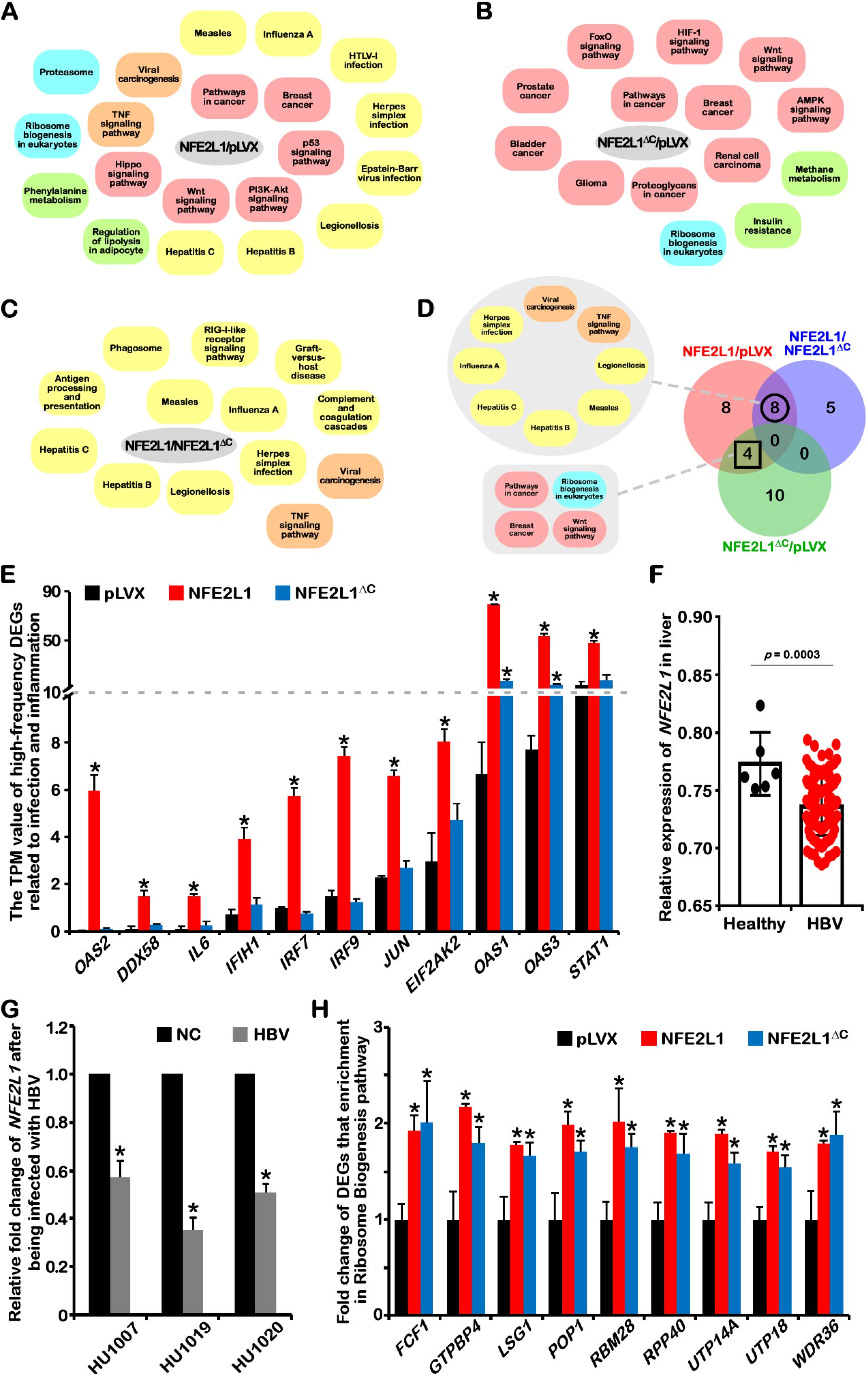
The enrichment of DEGs in KEGG pathway shows the features of NFE2L1’s transcription activator function and non-transcription factor activity. (A) Enrichment result of DEGs from the NFE2L1/pLVX group in KEGG pathway. (B) Enrichment result of DEGs from the NFE2L1^ΔC^/pLVX group in KEGG pathway. (C) The enrichment result of DEGs from NFE2L1/NFE2L1^ΔC^ group in KEGG pathway. (D) The Venn diagram was used to analyze the enrichment results of the three groups of DEGs to distinguish the transcription factor function and non-transcription factor function of NFE2L1. (E) Transcriptome data shows the expression of high-frequency DEGs (*OAS2*, *DDX58*, *IL6*, *IFIH1*, *IRF7*, *JUN*, *EIF2AK2*, *OAS1*, *OAS3*, *STAT1*) that involved in infection and inflammation. (F) The expression of *NFE2L1* gene in the livers of patients with HBV infection or healthy people tissues. The data comes from GEO (GSE83148) and *β-actin* was used as the internal reference gene to normalize the expression value of the target gene, red dots means the livers were HBV infected (HBV, 122 samples), and black dots means healthy livers (Healthy, 6 samples). (G) The effect of HBV infection on *NFE2L1* gene expression. Data from GEO (GSE112118) and *β-actin* was used as the internal reference. HU1007, HU1019, HU1020 are donor’s number. The “NC” group is a normal control group without treatment for 28 days, while the "HBV" group is an experimental group with HBV infection singly (HU1007) or co-infection with HDV (HU1019, HU1020) for 28 days. (H)The fold expression of DEGs (*FCF1*, *GTPBP4*, *LSG1*, *POP1*, *RBM28*, *RPP40*, *UTP14A*, *UTP18*, *WDR36*) that enrichment in Ribosome Biosynthesis in NFE2L1 and NFE2L1^ΔC^ cell lines compared with pLVX cell lines. n ≥ 3, ‘**\***’ means compared with pLVX group and *p* < 0.05.

Fourteen KEGG pathways with *p*-value < 0.01 obtained from the enrichment of DEGs in NFE2L1^ΔC^/pLVX group. Eleven of them were involved in development, cancer, and canceration (Wnt signaling pathway, FoxO signaling pathway, Renal cell carcinoma, HIF-1 signaling pathway, Breast cancer, Proteoglycans in cancer, Prostate cancer, Bladder cancer, Glioma, Pathways in cancer, AMPK signaling pathway). Two were related to metabolism (Methane metabolism, Insulin resistance), and one was Ribosome biogenesis in eukaryotes (Figs.4B and S1E).

To highlight the function of NFE2L1’s transcription factors, the NFE2L1 group data was compared with that of NFE2L1^ΔC^ group. The results showed that there were thirteen KEGG pathways with *p*-value < 0.01 obtained from the enrichment of DEGs in the NFE2L1/NFE2L1^ΔC^ group. Surprisingly, these pathways (Herpes simplex infection, Influenza A, Measles, Hepatitis C, Antigen processing and presentation, Complement and coagulation cascades, Graft-versus-host disease, RIG-I-like receptor signaling pathway, TNF signaling pathway, Hepatitis B, Phagosome, Legionellosis, Viral carcinogenesis) were all relevant to infection, inflammation and immunity (Figs.4C and S1F).

Then, the Venn diagram analysis was conducted based on the results of the above three groups of KEGG pathways. The results displayed eight pathways (Measles, Herpes simplex infection, Influenza A, Legionellosis, Viral carcinogenesis, TNF signaling pathway, Hepatitis C, Hepatitis B) intersecting the NFE2L1/pLVX group and the NFE2L1/NFE2L1^ΔC^ group were all correlated with infection and inflammation, while the four pathways (Ribosome biogenesis in eukaryotes, Pathways in cancer, Wnt signaling pathway, Breast cancer) intersecting the NFE2L1/pLVX group and the NFE2L1^ΔC^ /pLVX group were mainly related to development and cancer (Fig. 4D).

Some high-frequency DEGs (*JUN*, *STAT1*, *OAS1*, *OAS2*, *OAS3*, *IL6*, *IFIH1*, *IRF7*, *IRF9*, *EIF2AK2*, *DDX58*) that are involved in the infection and inflammation process were screened out. The results revealed that up-regulation of these gene caused by NFE2L1 strictly depended on its transcription factor function, in consideration of that NFE2L1^ΔC^ overexpression had no significant effects on most of those genes (Fig.4E).

HBV infection, one of the major global health problems, is the direct cause of hepatitis B and can elicit cirrhosis and HCC[32]. Specific knockout of NFE2L1 in the liver tissue of adult mice induced spontaneous non-alcoholic steatohepatitis (NASH) and HCC[9]. Combined with the results of transcriptome sequencing, NFE2L1 has been involved in infection and inflammation and carcinogenesis, suggesting the crucial position of NFE2L1 in both pathological processes of viral infection of the liver and the occurrence of further canceration.

To explore the correlation between NFE2L1 and virus infection, the relevant data from the Gene Expression Omnibus (GEO) database was analyzed. The results revealed that the expression of *NFE2L1* was significantly down-regulated in 122 hepatitis tissues infected with HBV compared with 6 normal liver tissues (Fig.4F). Another experiment with HBV infection of primary human hepatocytes also showed that after 28 days of HBV infection, the expression of NFE2L1 gene in liver cells was significantly suppressed (Fig.4G).

These data were in line with the transcriptome sequencing results and also indicated that the NFE2L1 may have a physiological function against infection by exogenous pathogenic microorganisms. Obviously, the inhibition of *NFE2L1* gene by HBV may be related to the HCC caused by HBV infection.

On the other hand, it is worth noting that the changes of ‘Ribosome biogenesis in eukaryotes’ pathway is most significant in responding to NFE2L1 function. Both NFE2L1 and NFE2L1^ΔC^ could affect the expression of DEGs (*FCF1*, *GTPBP4*, *LSG1*, *POP1*, *RBM28*, *RPP40*, *UTP14A*, *UTP18*, *WDR36*) in ‘Ribosome biogenesis in eukaryotes’ pathway, demonstrating that these genes are regulated by NFE2L1 in a non-transcription factor manner (Fig.4H). Combing with the classical regulation of NFE2L1 on Proteasome, we proposed the hypothesis that “the different subcellular localization of NFE2L1 may directly affect the protein synthesis and degradation process”, which means that NFE2L1 has an irreplaceable regulatory role in protein turnover.

Based on the previous analysis of DEGs enrichment in KEGG pathway, we could conclude that among the multiple molecular and physiological functions of NFE2L1, the regulation of redox homeostasis[18,33], proteasome[30,31], infection and inflammation mainly depends on its transcription activator function, and the regulation of metabolism, ribosome function, development and canceration are more related to its non-transcription factor activity.

### The tumor suppressive effect of NFE2L1 does not depend on its transcription factor function

Our previous studies confirmed that NFE2L1 could act as a tumor suppressor[10,11]. Here, the transcriptome sequencing results indicated that the regulation of NFE2L1 on development and canceration has been associated with its non-transcription factor activity (Fig. 4). So, we tested whether NFE2L1’s tumor suppressor effect depends on its transcription factor activity by performing subcutaneous tumor-bearing experiments in nude mice.

Compared to the control group (pLVX group), both the NFE2L1 and NFE2L1^ΔC^ overexpressed HepG2 cell lines exhibited slower growth rate, smaller tumor size, and lighter weight in nude mice bearing subcutaneous tumors without significant differences in overall body weight (Fig.5A-D). Consistently, when NFE2L1 was knocked down, the subcutaneous tumor grew dramatically (Fig. S3A-B). These results confirmed the tumor suppressive function of NFE2L1. And it was the first time to identify the tumor-suppressing effect of NFE2L1^ΔC^ without transcription factor activity, which proved that the tumor suppressive effect of NFE2L1 does not depend on its transcription factor function.

On the other hand, in the NFE2L1 group, tumor regressed in two of the five mice, while with one in the NFE2L1^ΔC^ group (Fig.5C). The tumor volume and weight of the NFE2L1 group are smaller and lighter than the NFE2L1^ΔC^ group’s (Fig.5C and D), means that the transcription activator function of NFE2L1 could enhance its tumor suppressive effect.

### NFE2L1 inhibits the Wnt/β-catenin signaling pathway

To explore the molecular mechanism of NFE2L1’s tumor suppression function, the DEGs that enriched in development- and canceration-related pathways were counted. 12 and 7 high-frequency DEGs were obtained from the NFE2L1/pLVX and NFE2L1^ΔC^/pLVX groups, respectively. Then intersection of these two sets of high-frequency DEGs yields five common high-frequency genes, namely *CCND1*, *CDKN1A*, *FZD4*, *FZD8*, and *WNT6* (Fig.5E).

Excluding cell cycle-related genes *CCND1* and *CDKN1A*, the *FZD4*, *FZD8* and *WNT6* all belong to Wnt signaling pathway, which plays crucial role in embryogenesis, organogenesis and homeostasis and can be activated in many types of cancer[34,35]. The expression of *WNT6* gene was decreased after overexpression of NFE2L1 or NFE2L1^ΔC^. However, *FZD4* and *FZD8,* genes encoding WNT6’s receptors[36], were increased (Fig. 5F). In addition, the transcriptome data showed that NFE2L1 overexpression significantly increase *FZD10* and decrease *WNT7B* and *WNT9A*. Meanwhile, the expression level of *WNT9A* was decreased and *WNT10B* was increased in NFE2L1^ΔC^ overexpressed cell lines (Fig. S2).

**Fig. 5.**
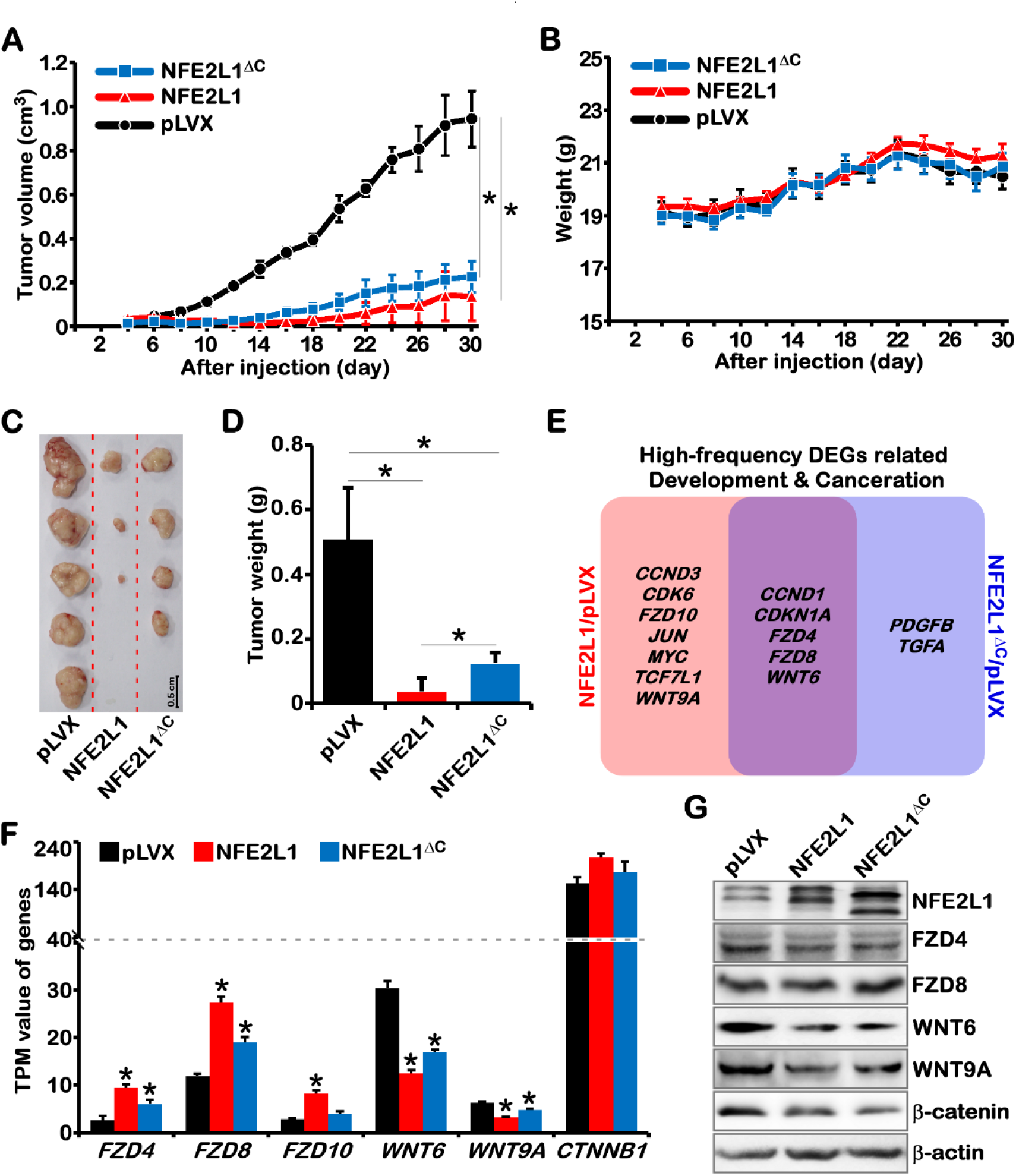
Both NFE2L1 and NFE2L1^ΔC^ can suppress tumors. 1×10^6^ cells/mouse experimental cells (pLVX, NFE2L1, and NFE2L1^ΔC^) were subcutaneously inoculated into nude mice (n = 5), the tumor volume (A) and body weight (B) were measured every two days. Photographs (C) and weight statistics (D) of tumors on the 30 th day. n ≥ 3, “**\***” means *p* < 0.05, and the scale is 0.5 cm. (E) The results of screening high-frequency DEGs, which come from NFE2L1/pLVX group and NFE2L1^ΔC^/pLVX group data sets and enrichment in development- and canceration-related pathways. (F) The expression of *FZD4*, *FZD8*, *FZD10*, *WNT6*, *WNT9A*, *CTNNB1* in three cell lines, the data comes from transcriptome sequencing. n = 3, “**\***” means compared with pLVX group and *p* < 0.05. (G) The changes of NFE2L1, FZD8, WNT6, WNT9A, β-catenin (CTNNB1) and β-actin protein in three cell lines were detected by WB.

Although the expression of *FZDs* were increased, the decrease of *WNTs* indicates that the Wnt signaling pathway may be suppressed from the source. On the other hand, Western Blotting results demonstrated that WNT6, WNT9A and β-catenin protein levels were reduced in NFE2L1 and NFE2L1^ΔC^ overexpressed cell lines, which confirmed the inhibition of Wnt/β-catenin signaling pathway by NFE2L1 (Fig. 5G). And in the shNFE2L1 cell line, we also detected over-activated β-catenin (Fig. S3C). Moreover, this result is consistent with the data of our previous discovery that NFE2L1 knockout resulted in an increase in β-catenin protein[37]. The Wnt/β-catenin pathway is the most frequently deregulated pathway in HCC, here we further found that the inhibition of NFE2L1 on Wnt/β-catenin pathway does not depend on its transcriptional activator function, this may be the molecular mechanism of HCC caused by *NFE2L1* deletion.

### NFE2L1 regulates the cell cycle through its non-transcription factor activity

Besides the Wnt signaling pathway, the regulation of the cell cycle may also be an important factor for NFE2L1 to inhibit tumor growth (Fig. 5E). By performing heatmap analysis on the cell cycle related genes that detected in the transcriptome sequencing, it was found that the effect of NFE2L1^ΔC^ on cell cycle related genes is basically the same as that of NFE2L1 (Fig. 6A). Through analyzing the key genes involved in regulating the multiple phase of cell cycle, the *CDKN2C*/*p18*, *CDKN2D*/*p19*, *CCND1*/*cyclin D1*, *CCND3*/*cyclin D3*, *CDKN1A*/*p21*, *CDKN1C*/*p57* and *CDC25A* could be affected by NFE2L1 and NFE2L1^ΔC^ (Fig. 6B).

**Fig. 6.**
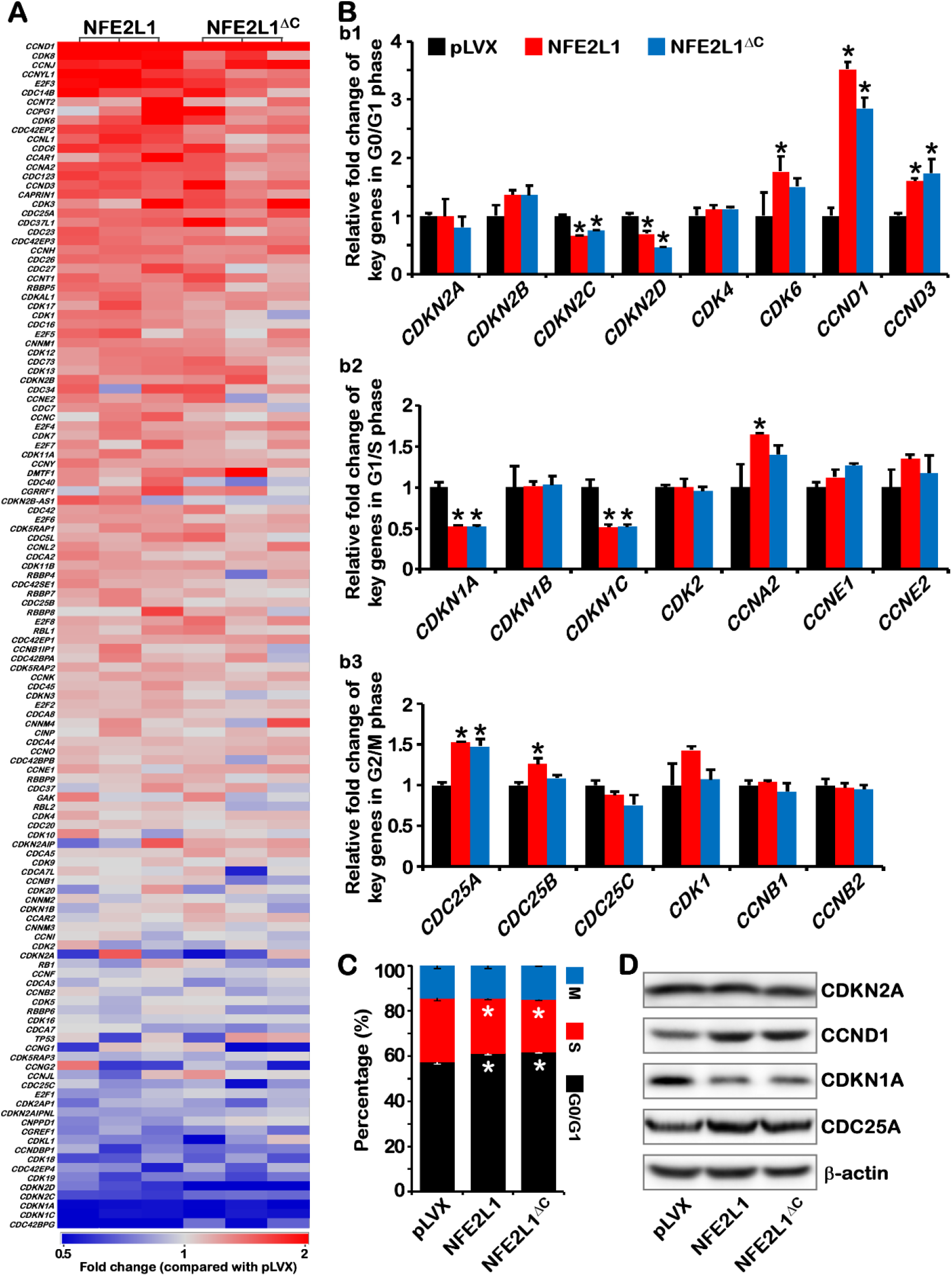
Regulation of cell cycle related genes by NFE2L1 and NFE2L1^ΔC^. (A) The heat map shows the expression of cell cycle related genes after overexpression of NFE2L1 or NFEL1^ΔC^ (red indicates an increase, blue indicates a decrease, and the data comes from transcriptome sequencing). (B) The effect of NFE2L1 or NFEL1^ΔC^ on the key regulatory genes in G0/G1 phases cell cycle arrest (b1), G1/S phases cell cycle arrest (b2), and G2/M phases cell cycle arrest (b3). (C)Flow cytometry analysis of the cell cycle of pLVX, NFE2L1 and NFE2L1^ΔC^ cell lines. (D) The changes of CDC25A, CCND1, CDKN1A, CDKN2A and Tubulin protein in three cell lines were detected by WB. n ≥ 3, ‘**\***’ means compared with pLVX group and *p* < 0.05.

The inhibition of *CDKN2C/p18*, *CDKN2D/p19* and the activation of *CCND1/cyclin D1*, *CCND3/cyclin D3* (Fig. 6b1) hinted that both NFE2L1 and NFE2L1^ΔC^ could promote cells to pass G0/G1 phase checkpoint[38]. *CDKN1A*/*p21* and *CDKN1C*/*p57*, the key regulators result in cell cycle arrest at G1/S phase[38], were suppressed both by NFE2L1 and NFE2L1^ΔC^ (Fig. 6b2), suggesting that NFE2L1 and NFE2L1^ΔC^ contribute to G1/S phase cell cycle progression. *CDC25A*, the key regulator that promotes G2/M transition[39], could be significantly activated by NFE2L1 and NFE2L1^ΔC^ (Fig. 6b3). Furthermore, both NFE2L1 and NFE2L1^ΔC^ could increase the expression of *E2F3* and *E2F4* and had no significant effects on the expression of *RB1* and *TP53* (Fig. S4A).

Transcriptome data indicated that overexpression of NFE2L1 or NFE2L1^ΔC^ induced similar effects on cell cycle. Flow cytometry results also showed that both overexpression of NFE2L1 or NFE2L1^ΔC^ increased the proportion of cells in the G0/G1 phase and decreased the ratio of cells in the S phase (Figs. 6C and S4B). Then, the changes in CDKN2A, CCND1, CDKN1A and CDC25A proteins were detected and the results were consistent with changes in their mRNA levels (Fig. 6D). These results revealed that although cell cycle-related genes tend to promote cell cycle progression, NFE2L1 induced cell cycle arrest in an unknown manner which has been independent of its transcription factor function.

## DISCUSSION

The transcription factor activity of NFE2L1 has been clearly described, which is achieved by forming a heterodimer with sMAF protein to recognize ARE elements [24]. In this article, we verified that NFE2L1 could act other than transcription activator by constructing a truncated protein NFE2L1^ΔC^ with no transcription factor activity (Figs. 1–3). The transcriptional repression function of NFE2L1 is germanely to its non-transcription factor function. This may be the reason that NFE2L1, as a transcription activator, exhibits transcription repression on some genes (such as *PTGS1*[14], *CD36*[23], *PPARγ*[40]). And almost all DEGs in the NFE2L1/NFE2L1^ΔC^ group were upregulated, which indicated the validity to characterize the transcription factor function of NFE2L1. However, 14 DEGs were still reduced (Table S1). Our previous research reported that NFE2L1 inhibited the expression of *PTEN* by directly up-regulating *mir-22*[14], suggesting that those 14 genes may be negatively regulated by NFE2L1 indirectly. These phenomena manifest that NFE2L1 has polymorphisms in the regulation of downstream genes, directly or indirectly.

Surprisingly, the transcriptome results revealed the dual regulation of NFE2L1 on ribosome and proteasome (Figs. 3, 4 and S1D), which depicted a spatio-temporal characteristic of NFE2L1’s function on protein turnover. NFE2L1 promoted protein synthesis by enhancing ribosomal function when it was stored in the cytoplasm and accelerated the degradation of ubiquitinated protein by improving the function of the UPS system when it was stimulated and transferred into the nucleus. The important role of NFE2L1 in the regulation of cellular redox homeostasis hints that NFE2L1 functions as an important factor to mediate the interaction between redox homeostasis and protein homeostasis in life system.

The transcriptome data displayed that NFE2L1 transcription factor activity was involved in the regulation of infection and inflammation (Fig. 4A, C and D) and the impact of viral infection on NFE2L1 expression was analyzed with the GEO database (Fig. 4F and G). These results implied that NFE2L1 may be an important transcription factor which endogenously activates the antiviral infection response. Additionally, the data implied that the tumor suppressive effect of NFE2L1 might be achieved by blocking the Wnt/β-catenin signaling pathway and inducing cell cycle arrest (Fig. 5, 6 and S3), suggesting the crucial molecular function of NFE2L1 in the cytoplasm. The inhibitory effect of NFE2L1 on Wnt signaling is consistent with its phenotype during embryonic development[16,17]. Data showing that NFE2L1 can inhibit Wnt/β-catenin signaling through its transcription factor-dependent activation of the Proteasome[37], this may be the reason for full-length NFE2L1 has stronger tumor suppressor activity (Fig. 5C and D).

In addition, HBV/HCV infection activates Wnt/β-catenin signaling[41,42] and inhibiting the expression of NFE2L1 that inhibits Wnt/β-catenin pathway, which suggests that NFE2L1 may be a key molecule that mediates HBV/HCV infection and HCC. However, the directly mechanism of NFE2L1’s inhibitory effect on Wnt/β-catenin pathway in the cytoplasm is still unclear. The decreased of β-catenin protein (Fig. 5G) suggest that the expressions of *FZD4*, *FZD8* and *FZD10* were increased due to NFE2L1 overexpression may be a compensation for the diminished of Wnt signaling. The phenotype of *NFE2L1* gene-edited animals and the effect of NFE2L1 on the WNT/β-catenin signaling pathway and cell cycle were consistent with the reported function of AXIN1 which is an important inhibitor for WNT/β-catenin signaling[43,44]. Studies have also shown that the AXIN1 protein can directly bind to the Neh4 (Nrf2– ECH homology 4) and Neh5 regions of the NFE2L2 protein[45]. NFE2L1 is a transcription factor of the same family of NFE2L2, with similar structural regions Neh4L (Nrf2–ECH homology 4 like) and Neh5L to Neh4 and Neh5[2]. All these evidences suggest that the anti-tumor effect mediated by NFE2L1 in the cytoplasm may be directly related to AXIN1.

Furthermore, NFE2L1 is also an important molecule for ER function to regulate ER stress[46], which means that NFE2L1’s non-transcription factor function may closely relate to ER function. Studies have shown that the ER membrane sensor NFE2L1, which directly binds and specifically senses cholesterol in the ER by its cholesterol recognition amino acid consensus (CRAC) domains, is central to cholesterol homeostasis[23,47]. This suggests that the non-transcription factor function of NFE2L1 may be related to cholesterol homeostasis in cells.

According to the research on the function of NFE2L1 in this study, we could conclude that besides the regulation of redox homeostasis and proteasome function, the transcriptional activation function of NFE2L1 has been implicated in mediating infection and inflammation-related effects (Fig. 7). The effects of NFE2L1 on metabolism, ribosome function, development and canceration are involve to its non-transcription factor activity (Fig. 7). NFE2L1 fulfills its non-transcription factor activity in the cytoplasm to promote cellular procession. And it translocates into the nucleus to perform transcription activator functions to enhance the ability of cells to resist environmental pressure when cells are stimulated.

**Fig. 7.**
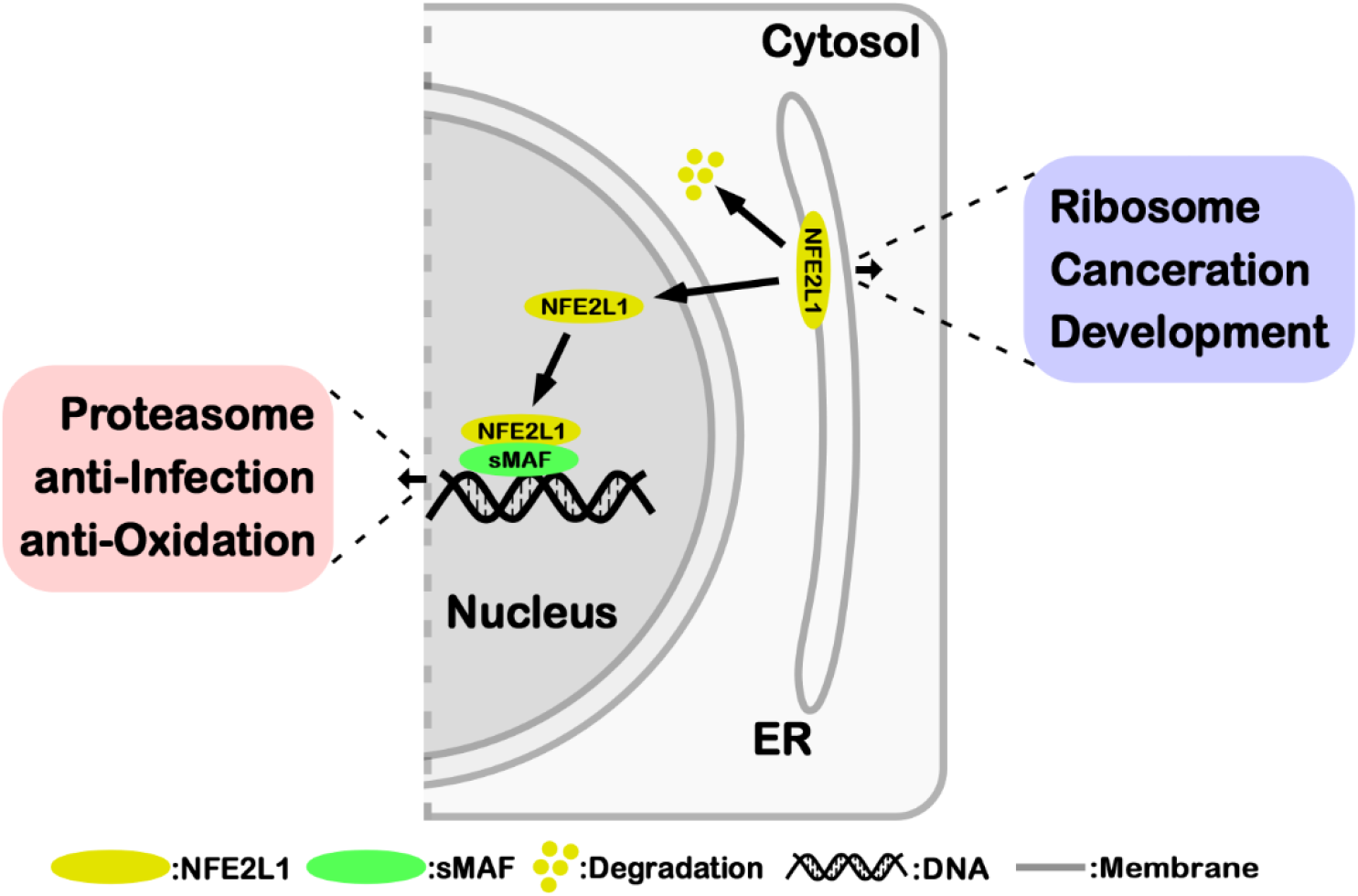
The model for the functional of NFE2L1 in nucleus and cytosol.

Transcription factors exhibit a wide range of molecular biological and physiological functions for the direct regulation of many downstream genes with different functions, making it difficult to utilize them as targets in clinical treatment studies. Our results firstly demonstrated that the tumor suppressive effect of NFE2L1 did not depend on its transcription factor activity, making it possible to target NFE2L1 for cancer treatment in cytoplasm without its transcription activator function in the nucleus.

In this article, we preliminarily revealed and described the non-transcription factor activity of NFE2L1. Further studies are needed to deeply understand this molecular, such as, the details how NFE2L1 mediates or executes its non-transcription factor activity in the cytoplasm, the precise process of regulating protein homeostasis through the dual role of ribosomes and proteasomes, the mechanism of NFE2L1 in tumor suppression and the relationship between NFE2L1 and WNTs in tumor suppression. In-depth knowledge will deepen the understanding of *NFE2L1* gene and lay foundations for targeting this gene for cancer treatment.

## MATERIALS AND METHODS

### Cell lines, culture and transfection

HEK293T cell line and human liver hepatocellular carcinoma cell line HepG2 were purchased from the Cell Bank of the Chinese Academy of Sciences (Shanghai), and cultured in DMEM medium (GIBCO, USA) in the 37°C incubator with 5% CO_2_. The DMEM medium was supplemented with 10% FBS (BI, Israel), 100 U/mL penicillin (Solarbio, China) and 100 μg/mL streptomycin (Solarbio, China). The cells were transfected with indicated plasmids alone or in combination for 8 h by using Lipofectamine 3000 (Thermo Fisher, USA), and then continue to cultivate for 24 h in the fresh medium before being subjected to indicated experiments.

### Constructs of plasmids

The NFE2L1 and NFE2L1^ΔC^ expression plasmid *pNFE2L1* and *pNFE2L1*^*ΔC*^ were made by cloning the target sequences from full-length CDS sequences of *NFE2L1* into *pLVX-Puro* vector. The luciferase plasmid *pGL3-ARE* which driven by the ARE element was obtained by inserting the 6×ARE sequences into the *pGL3-Basic* plasmid. The *pshNFE2L1* plasmid were constructed by *pLKO.1-puro* vector, and targeted at 5’- AAGATCTCATGTCCATCATGG-3’, 5’-AATCAGAATGTCAGCCTGCAT-3’ and 5’-GGACTTCTCCAGTTCACCA-3’, respectively. The sequences of the primers and oligos used to construct the plasmid are as follows: *NFE2L1-F*: 5’- CTCGAGGAATTCATGCTTTCTCTGAAGAAATAC-3’, *NFE2L1-R*: 5’- GGCCGCTCTAGATCACTTTCTCCGGTCCTTT-3’, *NFE2L1^ΔC^-R*: 5’- GAAAGGTCTAGATCAGGCTCGGGCTCGGTGCTCAT-3’, *6×ARE-F*: 5’- CTAGCAATGACTCAGCAAAATGACTCAGCAAAATGACTCAGCAAAATGACTCAGCAAAATGACTCAGCAAAATGACTCAGCAAA-3’, *6×ARE-R*: 5’- AGCTTTTGCTGAGTCATTTTGCTGAGTCATTTTGCTGAGTCATTTTGCTGAGTCATTTTGCTGAGTCATTTTGCTGAGTCATTG-3’, *shNFE2L1-1F*: 5’- CCGGAAGATCTCATGTCCATCATGGCTCGAGCCATGATGGACATGAGATC TTTTTTTG-3’, *shNFE2L1-1R*: 5’- AATTCAAAAAAAGATCTCATGTCCATCATGGCTCGAGCCATGATGGACATGAGATCTT-3, *shNFE2L1-2F*: 5’- CCGGAATCAGAATGTCAGCCTGCATCTCGAGATGCAGGCTGACATTCTGA TTTTTTTG-3, *shNFE2L1-2R*: 5’- AATTCAAAAAAATCAGAATGTCAGCCTGCATCTCGAGATGCAGGCTGACATTCTGATT-3’, *shNFE2L1-3F*: 5’-CCGGAAGGACTTCTCCAGTTCACCACTCGAGTGGTGAACTGGAGAAGTCCTTTTTTTG-3’, *shNFE2L1-3R* : 5’- AATTCAAAAAAAGGACTTCTCCAGTTCACCACTCGAGTGGTGAACTGGAGAAGTCCTT-3’. All primers and oligos used for construct plasmids were synthesized by Sangon Biotech (Shanghai, China).

### Luciferase reporter assay

Each well of 12-well plates were seeded 1.0×10^5^ cells, and cells were transfected by using Lipofectamine 3000 mixture with luciferase plasmids alone or plus overexpression plasmids when cell confluence reaches 80%, and the pRL-TK plasmid serves as an internal control for transfection efficiency. The ratio of expression plasmid: luciferase plasmid: pRL-TK plasmid is 5: 5: 1. The luciferase activity was measured by the dual-luciferase reporter assay system (E1910, Promega, USA).

### Western blotting (WB) and antibodies

Experimental cells were harvested by using the total protein extraction buffer (Tris 7.58 g/L, SDS 20 g/L, β-mercaptoethanol 20 mL/L, Glycerol 100 mL/L, Bromophenol Blue 0.2 g/L, pH6.8) and the lysates were denatured immediately at 100°C for 10 min. Subsequently, equal amounts of protein extracts were subjected to separation by SDS-PAGE, and subsequent visualization was performed by Western blotting with indicated antibodies. The monoclonal antibody of NFE2L1 (D5B10) was purchased from CST, USA. The antibody of Ubiquitin (ab134953), WNT6 (ab154144), CDKN1A (ab109520) and CDKN2A (ab81278) were purchased from Abcam, USA. The antibody of FZD8 (D261206), WNT9A (D121400), CDC25A (D155203), β-catenin (D260137) were purchased from Sangon Biotech, China. The antibody of CCND1 (GB11079) was purchased from Servicebio, China. The β-actin (GB11001, Servicebio, China) and Tubulin (ab210797, Abcam, USA) were served as internal control to correct the mass of samples.

### RNA isolation and real-time quantitative PCR (qPCR)

Experimental cells were subjected to isolation of total RNAs by using the RNAsimple Kit (DP419, Tiangen Biotech, China). Then, 1 μg of total RNA were added in a reverse-transcriptase reaction to generate the first strand of cDNA with the RevertAid First Strand Synthesis Kit (K1622, Thermo Fisher, USA). The synthesized cDNA was served as the template for qPCR and the GoTaq® qPCR Master Mix (A6001, Promega, USA) was used. The mRNA expression level of β-actin was served as an optimal internal standard control. The sequences of primers for qPCR: *HOMX1-F*: 5’- GCCAGCAACAAAGTGCAAGAT-3’, *HOMX1-R*: 5’-GGTAAGGAAGCCAGCCA AGAGA-3’, *PTGS1-F*: 5’-AGAAGGAGATGGCAGCAGAGT-3’, *PTGS1-R*: 5’-TAG GGACAGGTCTTGGTGTTGA-3’, *PTGS2-F*: 5’-GGTTGCTGGTGGTAGGAATGT T-3’, *PTGS2-R*: 5’-GCAGGATACAGCTCCACAGCAT-3’, *PPARGC1B-F*: 5’-TGGT GAGATTGAGGAGTGCGA-3’, *PPARGC1B-R*: 5’-GCCTTGTCTGAGGTATTGAGGTATTC-3’, *β-actin-F*: 5’-CATGTACGTTGCTATCCAGGC-3’, *β-actin-R*: 5’-CTCCTTAATGTCACGCACGAT-3’. All primers used for qPCR were synthesized by Sangon Biotech (Shanghai, China).

### Lentiviral packaging system and the construction of cell lines

Each well of 6 well plates were seeded 5×10^5^ HEK293T cells. The cells were transfected with 3 plasmids (1μg *pMD2.G*, 2μg *psPAX2*, or 3μg *pLKO.1*/*pshNFE2L1*/*pLVX-Puro*/*pNFE2L1*/*pNFE2L1*^*ΔC*^, and the transfection volume is 1mL) when cell confluence reaches 80%. The transfection reagent was replaced with fresh medium after transfection for 8 h and the medium was collected and filtered by using 0.45 μm sterile filter to obtain the virus after 36~48 h. The virus was used to transduce the target cells, puromycin was used to screen out the infected cells, and the overexpression or knockdown efficiency was verified by WB.

### Subcutaneous tumor xenografts in nude mice

All relevant animal experiments in this study were conducted according to the valid ethical regulations that have been approved. All mice were maintained under standard animal housing conditions. Mouse xenograft models were established by subcutaneously injected of the human hepatoma HepG2 (pLVX, NFE2L1, NFE2L1, shNC, or shNFE2L1). Experimental cells (1×10^6^) were allowed for growth in the exponential phase and then suspended in 0.2 mL PBS. Then, these cells were subcutaneously inoculated into male nude mice (BALB/c nu/nu, 7 weeks, HFK Bioscience, Beijing, China) at a single site. The tumor sizes were measured every two days. These mice were sacrificed and their transplanted tumors were excised on the 30th day.

### Cell cycle analysis

Equal numbers of pLVX, NFE2L1, NFE2L1^ΔC^cells were seeded in 6‐cm dish to reach 90% confluence. Then cells were collected and stained with a Cell Cycle Staining Kit (CCS012, Multi Sciences, China), and subsequently subjected to flow cytometry analysis.

### The transcriptomic RNA sequencing analysis

Total RNAs were subjected to the transcriptomic sequencing by the Sangon Biotech (Shanghai, China) on the HiseqNOVA platform (contract No.: MRNA193522ZZ). The clean reads were generated and mapped to the reference by using the HISAT2 tool[48] after removing the ‘dirty’ raw reads with data filtering. The gene expression levels were calculated by using the TPM (Transcripts Per Kilobase of exon model per Million mapped reads). Then, DEGs (differentially expressed genes) were identified with the criteria TPM > 5, |Fold Change| > 2, *p*-Value < 0.01 or TPM > 0.1, |Fold Change| > 1.5, *p*-Value < 0.01 by DESeq. The KAAS was used for KEGG annotation and cluster Profiler for KEGG enrichment analysis of DEGs[49,50].

### Statistical analysis

Significant differences were statistically determined by the Student’s *t*-test or Multiple Analysis of Variations (MANOVA). The data were shown as a fold change (means ± S.D.), each of which represents at least three independent experiments that performed in triplicate.

## Acknowledgement

This work was supported by Postdoctoral Research Grant in Henan Province (201901006) and National Natural Science Foundation of China (81822043 and U1604286).

## Supplementary Figures and Figure Legend

**Fig. S1.**
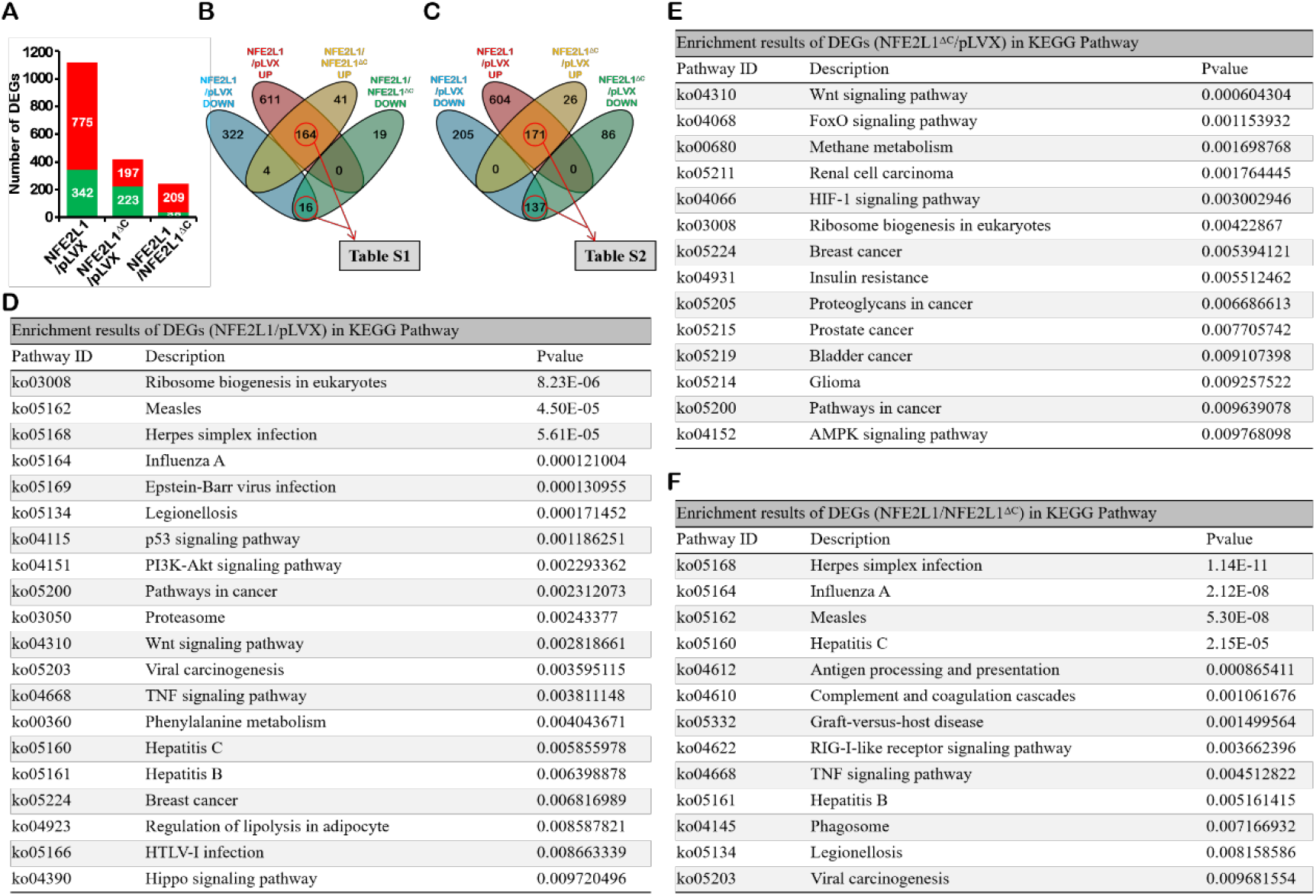
Screening, classification and functional enrichment of the downstream genes of NFE2L1. (A) Screening and counts of the DEGs in NFE2L1/pLVX, NFE2L1^ΔC^/pLVX, and NFE2L1/NFE2L1^ΔC^ under a new condition (TPM>0.1, |Fold Change|>1.5, *p*-value < 0.01), the red color means increased DEGs, the green color means decreased DEGs. (B) Screening the downstream genes that regulated by NFE2L1’s transcription activator function by intersection of DEGs that in NFE2L1/pLVX and NFE2L1/NFE2L1^ΔC^ groups. (C) The DEGs related to NFE2L1’s non-transcription factor activity was obtained though intersection of the DEGs from NFE2L1/pLVX, NFE2L1^ΔC^/pLVX groups. ((D) Enrichment results of DEGs in the NFE2L1/pLVX group in KEGG pathway with *p*-value < 0.01. (E) Enrichment results of DEGs in the NFE2L1^ΔC^/pLVX group in KEGG pathway with *p*-value < 0.01. (F) Enrichment results of DEGs in the NFE2L1/ NFE2L1^ΔC^ group in KEGG pathway with *p*-value < 0.01.

**Fig. S2.**
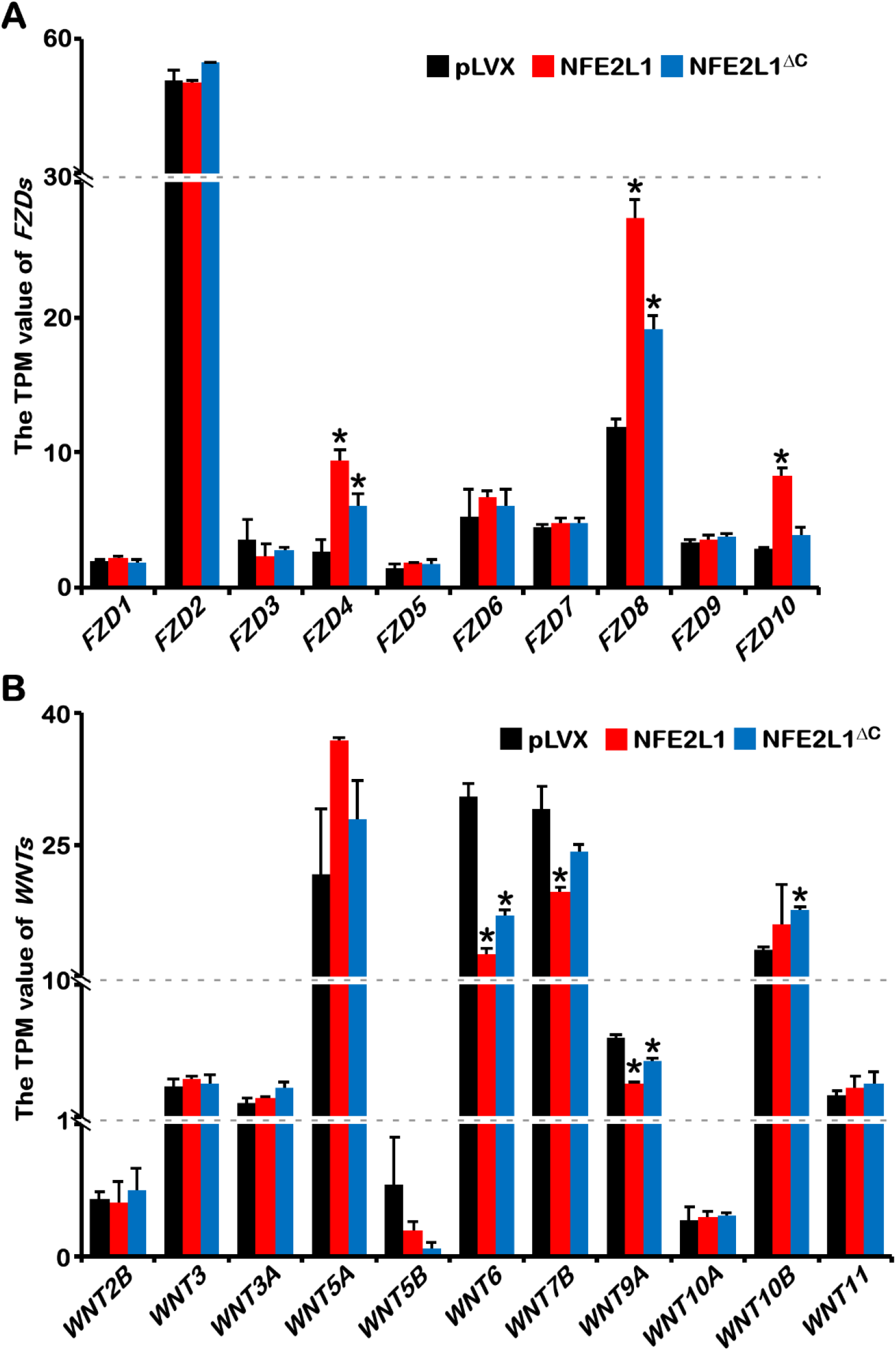
Transcriptome data presents the expression of *FZDs* and *WNTs* genes. (A) The effect of NFE2L1 and NFE2L1^ΔC^ overexpression on *FZD1*, *FZD2*, *FZD3*, *FZD4*, *FZD5*, *FZD6*, *FZD7*, *FZD8*, *FZD9* and *FZD10*. (B) The effects of NFE2L1 and NFE2L1^ΔC^ overexpression on *WNT2B*, *WNT3*, *WNT3A*, *WNT5A*, *WNT5B*, *WNT6*, *WNT7B*, *WNT9A*, *WNT10A*, *WNT10B* and *WNT11*. n=3, ‘**\***’ means compared with pLVX group and *p* < 0.05.

**Fig. S3.**
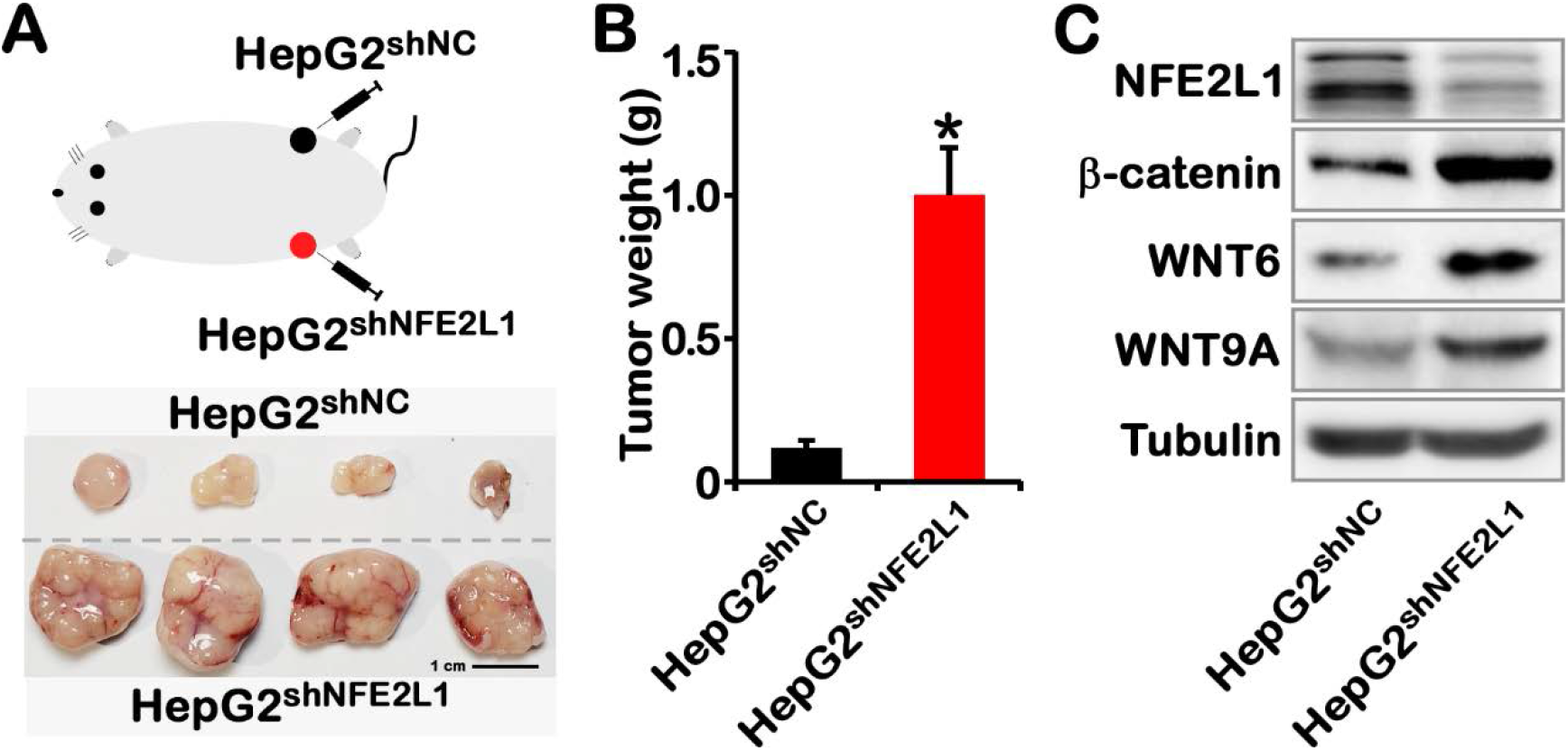
*NFE2L1* knockdown causes the malignant proliferation of HepG2 tumor-bearing tumors. The *NFE2L1* knockdown cell line HepG2^shNFE2L1^ and the control cell line HepG2^shNC^ were subcutaneously inoculated into nude mice (1×10^6^ cells each site), the tumors were dissected after 2 weeks, photographs (A) and weight statistics (B) of tumors. n=4, “**\***” means *p* < 0.05, and the scale is 1 cm. (C) The changes of NFE2L1, β-catenin, WNT6, WNT9A, Tubulin proteins in HepG2^shNC^ and HepG2^shNFE2L1^ cell lines were detected by WB.

**Fig. S4.**
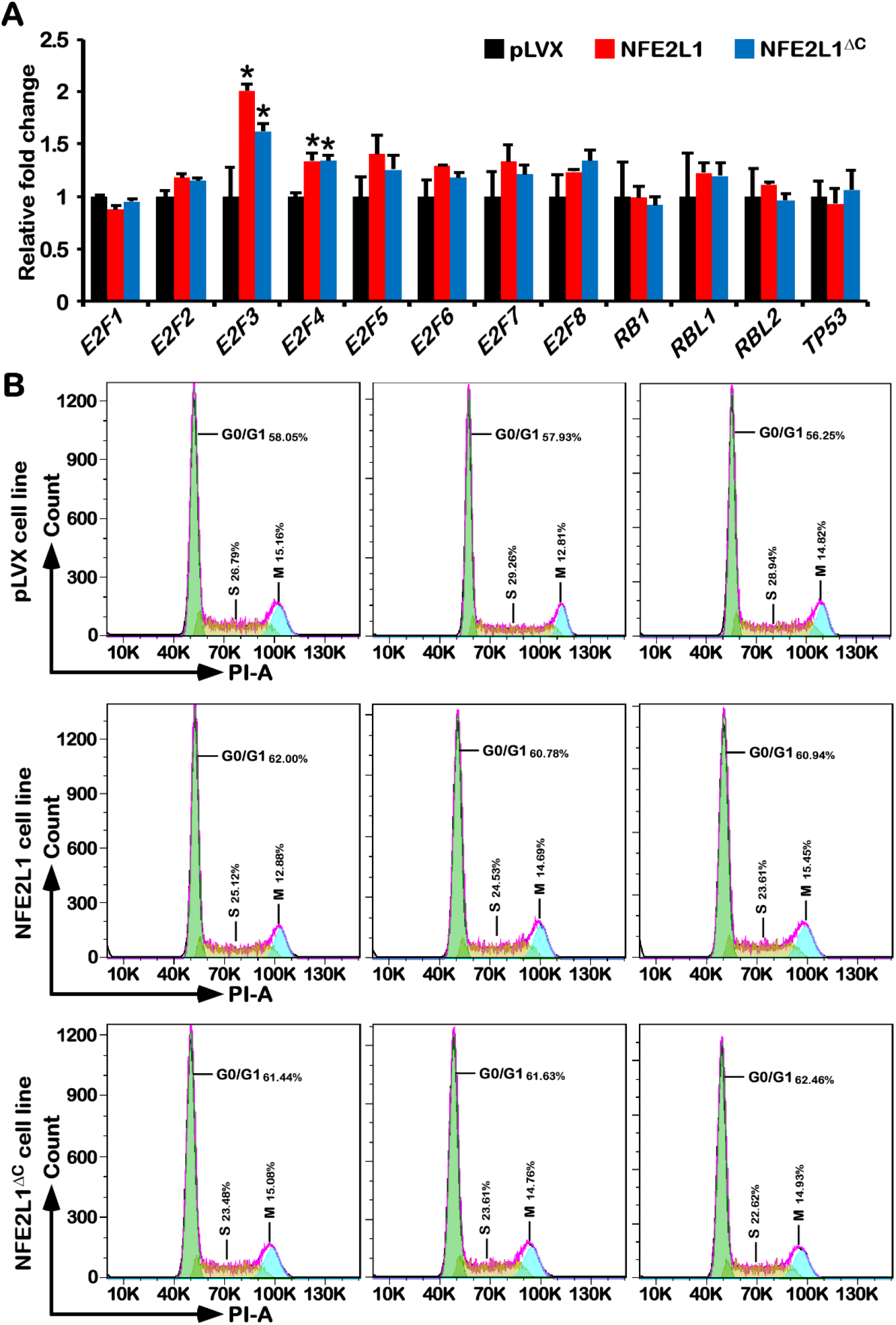
The effects of NFE2L1 and NFE2L1^ΔC^ on cell cycle. (A)The expression of cell cycle related genes (*E2Fs*, *RB1*, *RBL1*, *RBL2*, *TP53*) in pLVX, NFE2L1, NFEL1^ΔC^ cell lines. n≥3, ‘*’ means compared with pLVX group and *p* < 0.05. (B) Flow cytometry analysis of the cell cycle of pLVX, NFE2L1 and NFE2L1^ΔC^ cell lines.

## Supplementary Tables

**Table S1.**
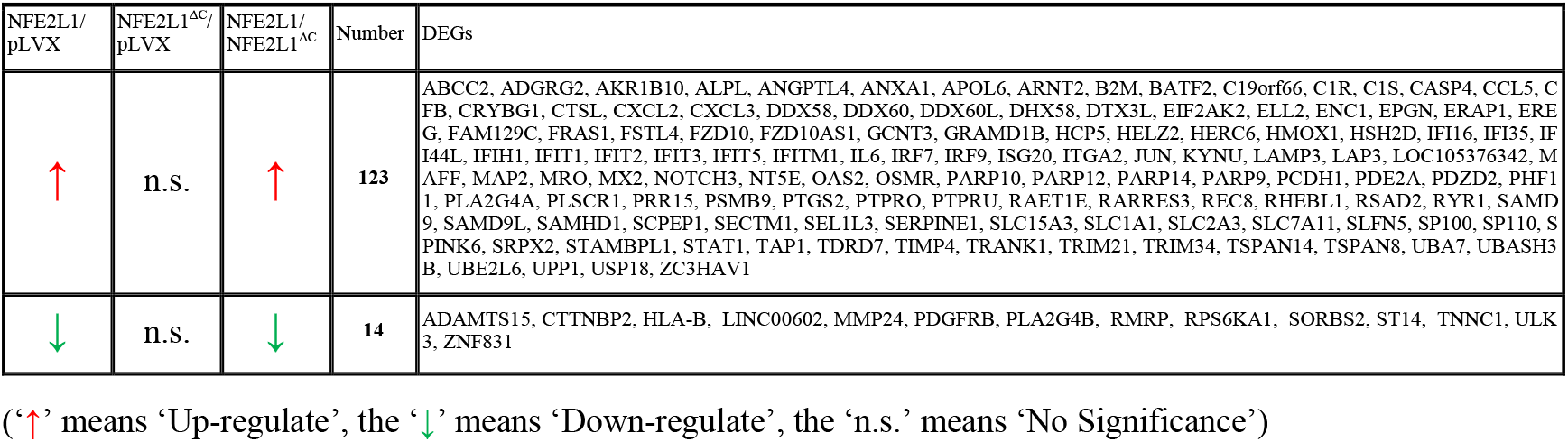
The DEGs that regulated by NFE2L1’s transcription activator function.

**Table S2.**
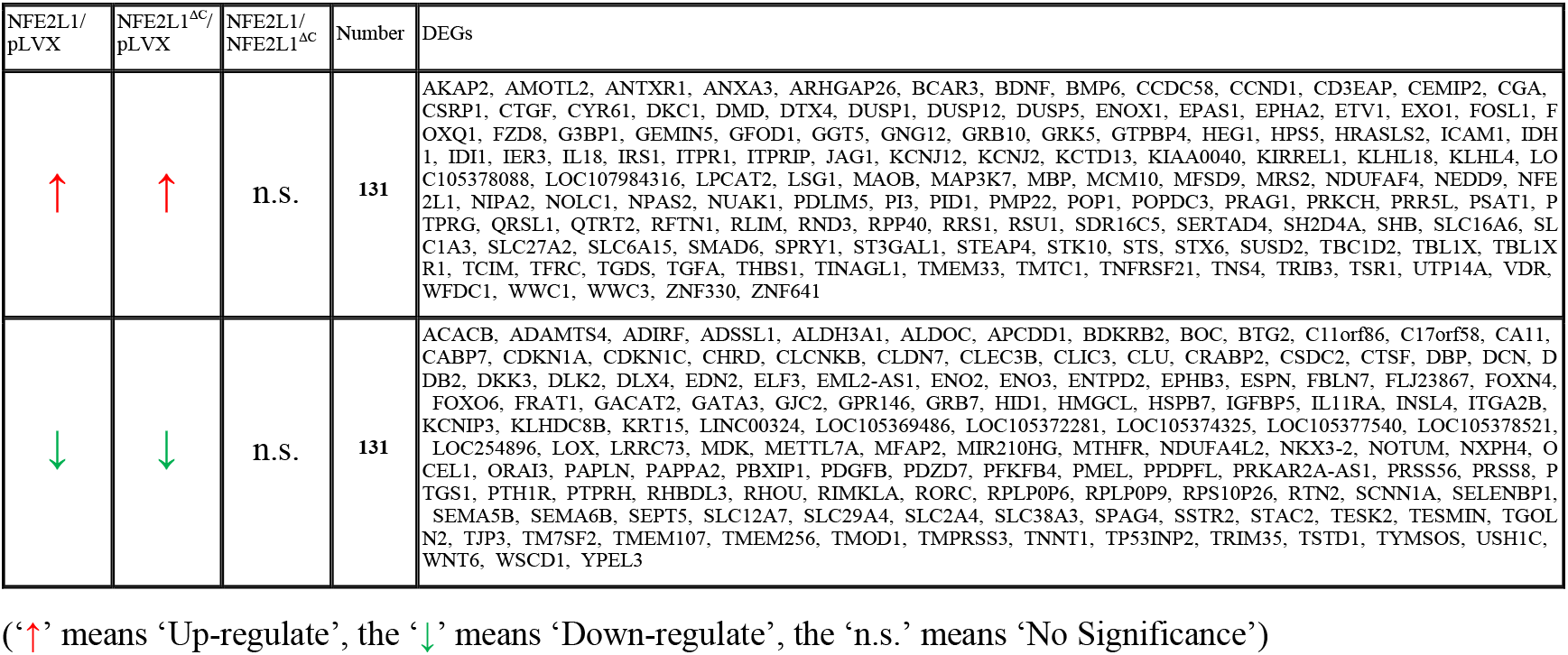
The DEGs that related to NFE2L1’s non-transcription factor activity.

